# Metabolite T_2_ relaxation times decrease across the adult lifespan in a large multi-site cohort

**DOI:** 10.1101/2024.06.19.599719

**Authors:** Kathleen E. Hupfeld, Saipavitra Murali-Manohar, Helge J. Zöllner, Yulu Song, Christopher W. Davies-Jenkins, Aaron T. Gudmundson, Dunja Simičić, Gizeaddis Simegn, Emily E. Carter, Steve C. N. Hui, Vivek Yedavalli, Georg Oeltzschner, Eric C. Porges, Richard A. E. Edden

## Abstract

**Purpose:** Relaxation correction is crucial for accurately estimating metabolite concentrations measured using *in vivo* magnetic resonance spectroscopy (MRS). However, the majority of MRS quantification routines assume that relaxation values remain constant across the lifespan, despite prior evidence of T_2_ changes with aging for multiple of the major metabolites. Here, we comprehensively investigate correlations between T_2_ and age in a large, multi-site cohort.

**Methods:** We recruited approximately 10 male and 10 female participants from each decade of life: 18-29, 30-39, 40-49, 50-59, and 60+ years old (*n*=101 total). We collected PRESS data at 8 TEs (30, 50, 74, 101, 135, 179, 241, and 350 ms) from voxels placed in white-matter-rich centrum semiovale (CSO) and gray-matter-rich posterior cingulate cortex (PCC). We quantified metabolite amplitudes using Osprey and fit exponential decay curves to estimate T_2_.

**Results:** Older age was correlated with shorter T_2_ for tNAA, tCr_3.0_, tCr_3.9_, tCho, Glx, and tissue water in CSO and PCC; r_s_ = −0.21 to −0.65, all *p*<0.05, FDR-corrected for multiple comparisons. These associations remained statistically significant when controlling for cortical atrophy. T_2_ values did not differ across the adult lifespan for mI. By region, T_2_ values were longer in the CSO for tNAA, tCr_3.0_, tCr_3.9_, Glx, and tissue water and longer in the PCC for tCho and mI.

**Conclusion:** These findings underscore the importance of considering metabolite T_2_ changes with aging in MRS quantification. We suggest that future 3T work utilize the equations presented here to estimate age-specific T_2_ values instead of relying on uniform default values.

## 1. Introduction

Understanding brain changes across the healthy adult lifespan is critical for preserving brain health in the quickly aging global population and uncovering possible mechanisms of age-related neurological disease. Proton magnetic resonance spectroscopy (^1^H MRS) is the only methodology that allows non-invasive measurements of endogenous brain metabolite concentrations. However, MRS data are often acquired at echo times (TEs) that are non-negligible compared to metabolite transverse relaxation rates (T_2_). This results in T_2_-weighting of the signal, such that metabolite amplitude changes associated with normal aging might be caused by changes in relaxation but misinterpreted as changes in metabolite concentration.

This confound exists even for short-TE MRS, but is particularly a concern for *J*-difference-edited MRS^1^, which relies on TEs of >65 ms (constrained by the duration of frequency-selective editing pulses and the J-evolution of target metabolites). Thus, recent consensus^1–3^ suggests that it is critical to address the confound of T_2_ relaxation (including for reference signals), particularly in studies of aging and neurodegeneration. However, despite this, most MRS quantification procedures (likely incorrectly) use static reference values, which assume that metabolite T_2_ relaxation remains constant across the adult lifespan.

Prior work has reported varied relationships between age and T_2_, primarily for the singlet resonances total *N*-acetyl aspartate (tNAA), creatine (tCr), and choline (tCho). A majority of prior work at 3 and 4 T has reported shorter metabolite T_2_s with older age both for metabolites^4–7^ and tissue water^4,7^. A few studies^8,9^ at 1.5 T have reported the opposite effect of longer metabolite T_2_s with older age; however, one of these studies^9^ was complicated by overlap among water and metabolite signals, and the other^8^ examined only the frontal lobe and included only males in the sample. One study^10^, also at 1.5 T, found longer NAA T_2_ in the centrum semiovale of older adults; however, this study utilized a linewidth-based approach which has not been validated or used again since publication in 2005. With the exception of work by Brooks and colleagues^8^, each of these prior studies involved comparison of discrete age groups (young versus older adults) rather than continuous sampling across the adult lifespan, and each used small sample sizes (all *n* < 20 per age group, with the exception of work by Deelchand and colleagues^4^ which included 32 young and 26 older adults). Therefore, in the present study, we leveraged a large, multi-site cohort in order to more comprehensively investigate whether metabolite and tissue water T_2_ values differ across the normal adult lifespan, and to provide statistical models for calculating age-specific T_2_ values for future integration into MRS quantification procedures.

## 2. Methods

### 2.1 Participants

101 healthy adults provided written informed consent to participate at one of two sites: the Johns Hopkins University School of Medicine (*n* = 51) and the University of Florida (*n* = 50). The sample included approximately 10 females and 10 males from each of the following decades: 18–29, 30–39, 40–49, 50–59, and 60+ years (Table 1). The Johns Hopkins University and University of Florida Institutional Review Boards approved all study procedures.

**Table 1.** Participant Demographics.

| Variable | 18-29 years | 30-39 years | 40-49 years | 50-59 years | 60+ years |
| --- | --- | --- | --- | --- | --- |
| Sex, # | 10 F, 13 M | 10 F, 10 M | 10 F, 10 M | 9 F, 9 M | 11 F, 9 M |
| Age, Mean (SD),<br>[Min, Max], years | 24.2 (3.3)<br>[18.7, 29.8] | 35.0 (2.9)<br>[30.3, 39.9] | 44.2 (3.0)<br>[40.0, 49.2] | 55.0 (2.9)<br>[50.2, 59.8] | 67.5 (4.1)<br>[60.8, 75.4] |
| MoCA, Mean (SD)<sup>a</sup><br>[Min, Max] | 27.9 (2.0)<br>[23, 30] | 28.0 (2.3)<br>[23, 30] | 27.6 (2.5)<br>[22, 30] | 26.3 (1.8)<br>[23, 29] | 27.7 (2.5)<br>[22, 30] |

<sup>a</sup> One point was added to the MoCA score for $n = 9$ , as these individuals reported having completed 12 years of education or less<sup>11</sup>.

Participants first completed the Montreal Cognitive Assessment (MoCA)^11^, followed by a 1-hour MRI protocol. Of note, four individuals scored below the cut-off score of 23 out of 30 (i.e., indicative of possible mild cognitive impairment^12^). However, MoCA score was not an *a priori* exclusion criterion for this study. Moreover, each of these individuals scored 22 (just below the cut-off), and 3 of these 4 reported that English was not their primary language which can negatively impact MoCA performance^13^ (and it was not feasible to conduct the MoCA in a language other than English). Therefore, we presumed that cognitive impairment was not likely and opted to retain these individuals in the cohort and statistical analyses.

### 2.2 MRS Acquisition

All scans were performed using a 32-channel head coil on either the Johns Hopkins University 3 T Philips dStream Ingenia Elition MRI scanner or the University of Florida 3 T Philips MR7700 MRI scanner. For voxel positioning, we first collected a *T*_1_-weighted structural MRI scan using the following parameters: MPRAGE, TR/TE 2000 ms/2 ms, flip angle 8°, slice thickness 1.0 mm, 150 slices, voxel size 1 mm^3^ isotropic, total time 2 min 46 sec. Next, we acquired TE series data from two 30 x 26 x 26 mm^3^ voxels: the white matter (WM) rich centrum semiovale (CSO) and the gray matter (GM) rich posterior cingulate cortex (PCC; Figure 1).

**Figure 1.**
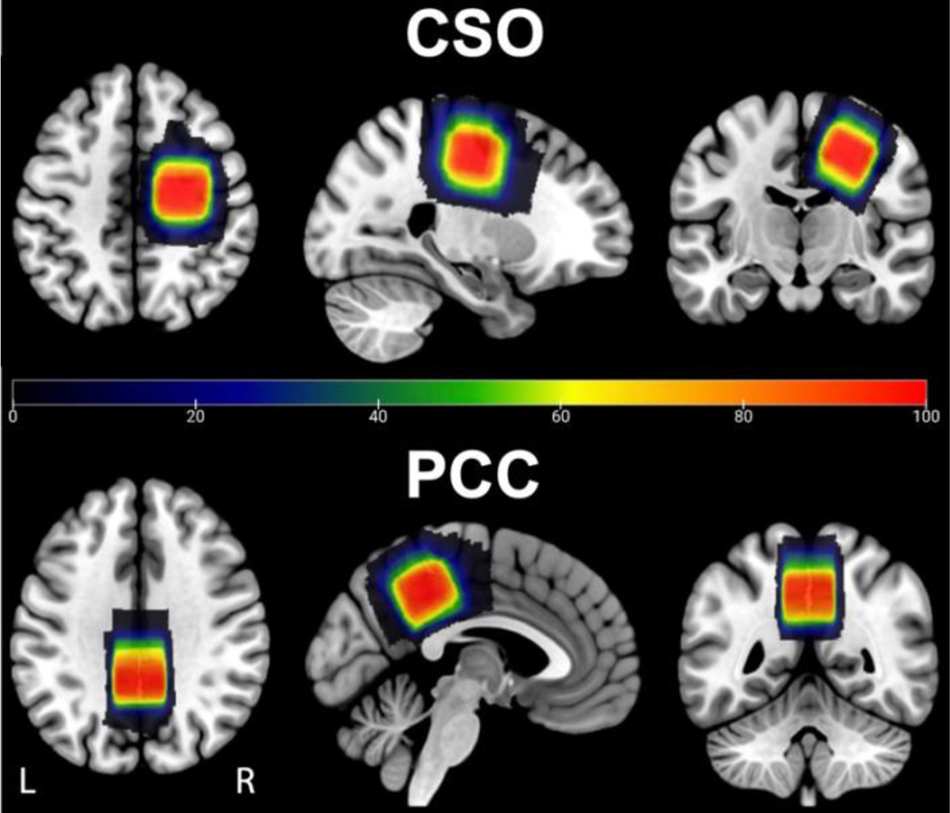
Voxel Placement. TE series data were acquired from the centrum semiovale (CSO) and posterior cingulate cortex (PCC). Each participant’s native space binary voxel mask for their CSO and PCC voxels was normalized to standard (MNI) space and overlaid onto the spm152 template. Warmer colors indicate areas of greater overlap between participants (color bar = number of subjects overlapped).

Scan parameters for the TE series included: PRESS localization, TR 2000 ms, 8 logarithmically-spaced TEs 30, 50, 74, 101, 135, 179, 241, and 350 ms, 24 transients per TE sampled at 2000 Hz with 1024 points, and CHESS water suppression (115 Hz bandwidth). Within each series, the TE steps were neither interleaved nor randomized. We also collected a separate series of unsuppressed water reference data at each of the 8 TEs with the same parameters, but with 2 transients per TE and no water suppression. Of note, these voxel sizes and locations were selected to match those collected in our recent cohort of short-TE PRESS metabolite data in 102 individuals ranging from their 20s to their 60s^14^.

### 2.2 MRS Data Processing

MRS data were analyzed within MATLAB R2021b using the open-source analysis toolbox Osprey (v2.5.0; https://github.com/schorschinho/osprey/)^15^. All analysis procedures followed consensus-recommended guidelines^1,3^. Briefly, analysis steps included: loading the vendor-native raw data (which had already been coil-combined, eddy-current-corrected, and averaged on the scanner at the time of data collection), removing the residual water signal using a Hankel singular value decomposition (HSVD) filter^16^, and modeling the metabolite peaks at each TE separately as described previously^15,17^ using TE-specific custom basis sets. The basis sets were simulated by the MRSCloud tool^18^ (https://braingps.mricloud.org/mrs-cloud).

MRSCloud using a localized 2D density-matrix simulation of a 101 x 101 spatial grid (voxel size 30 x 30 x 30 mm^3^; field of view 45 x 45 x 45 mm^3^) and vendor-specific refocusing pulse shape, duration, and sequence timings based on the MATLAB simulation toolbox FID-A^19^. The basis sets consisted of 18 basis functions: ascorbate (Asc), aspartate (Asp), creatine (Cr), negative creatine methylene (-CrCH_2_), gamma-aminobutyric acid (GABA), glycerophosphocholine (GPC), glutathione (GSH), glutamine (Gln), glutamate (Glu), lactate (Lac), myo-inositol (mI), *N*-acetyl aspartate (NAA), *N*-acetyl aspartyl glutamate (NAAG), phosphocholine (PCh), phosphocreatine (PCr), phosphoethanolamine (PE), scyllo-inositol (sI), and taurine (Tau), as well as 5 macromolecule signals (MM09, MM12, MM14, MM17, MM20) and 3 lipid signals (Lip09, Lip13, Lip20) included as parameterized Gaussian functions^17^.

We extracted amplitudes for 6 metabolites of interest: tNAA, tCho, tCr_3.0_ (Cr + PCr), tCr_3.9_ (Cr + PCr - (-CrCH_2_)), mI, and Glx (Glu + Gln). We multiplied each metabolite amplitude by Osprey’s internal *MRSCont.fit.scale* factor for each TE and participant to make the metabolite amplitudes directly comparable across TEs. This scaling factor is applied to the data to ensure an optimal dynamic range between the data and basis set during modeling. It is defined as the ratio of the maximum of the real part of the data and the basis set in the model range. Next, we used *lsqcurvefit* in MATLAB to fit monoexponential T_2_ decay functions to the TE series metabolite amplitudes (Equation 1) in order to obtain the T_2_ decay constant and an R^2^ value of model fit for each participant for each metabolite.

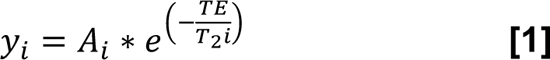

In Equation 1, y_i_ represents the metabolite amplitude, A_i_ is a scaling constant, and T_2i_ is the relaxation time to be calculated for the i^th^ subject. Lastly, we created a binary mask of the two MRS voxels in subject space, co-registered these masks to each participant’s T_1_-weighted structural scan, and segmented the structural scans using SPM12^20^, in order to calculate the volume fractions of white matter (fWM), gray matter (fGM), and cerebrospinal fluid (fCSF) in each participant’s voxels in subject space (for use in statistical models to control for cortical atrophy and for estimation of tissue water T_2_).

We then repeated a similar procedure for tissue water. To estimate the water amplitudes, the unsuppressed water data at each TE was modeled using a linear combination model with a simulated water signal^18^. We used MATLAB’s *lsqcurvefit* to fit a biexponential decay function (Equation 2) to obtain tissue water T_2_ and R^2^ values for each participant.

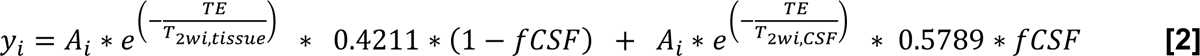

In Equation 2, y_i_ represents the water amplitude, A_i_ is a scaling constant, T_2wi,tissue_ is the tissue water relaxation time, and T_2wi,CSF_ is the CSF water relaxation time to be calculated for the i^th^ subject. fCSF is the fraction of CSF within the voxel for the i^th^ subject; 0.4211 weights the first term by the approximate molal concentration of water for non-CSF tissue (40/(40+55)), and 0.5789 weights the second term by the approximate molal concentration of water for CSF (55/(40+55)). The A_i_ and T_2wi,tissue_ terms were unconstrained, and the T_2wi,CSF_ term was constrained to the range of 50–3000 ms^21–23^. A data acquisition error occurred at the University of Florida site for the water data; therefore, we included only the Johns Hopkins University participants (*n* = 51) in statistical analyses of the water T_2_ data.

### 2.3 Statistical Analyses

We conducted all statistical analyses using R 4.3.2^24^ within RStudio^25^. First, we calculated descriptive statistics (mean, standard deviation) by age group for the T_2_ values for each of the 6 metabolites of interest and tissue water. Next, we examined the correlation between T_2_ and age for each metabolite and voxel separately. As multiple variables did not satisfy the Pearson correlation normality assumption (Shapiro test *p* < 0.05), we instead report nonparametric Spearman correlations. To account for multiple comparisons, we applied the Benjamini-Hochberg false discovery rate (FDR) correction to the *p*-values for each voxel^26^.

Secondly, we ran a series of linear models, setting each metabolite T_2_ as the outcome variable and age as the predictor: T_2_ = β_0_ + β_1_*(Age-30). We centered age around 30 years, so that the intercept (β_0_) from this model would represent the predicted metabolite T_2_ value at 30 years old, and the slope (β_1_) would represent the change in T_2_ for each year of age. The aim of this model was to provide an equation to calculate predicted T_2_ value for a given metabolite given the age of a participant. As a follow-up analysis, we reran each of these linear models controlling for the potential effects of cortical atrophy with aging: T_2_ = β_0_ + β_1_*(Age-30) + β_2_*Tissue. As in our recent work examining metabolite T_1_ changes with aging^27^, we calculated cortical atrophy as the relative tissue fraction within the voxel, fGM / (fWM + fGM). The purpose of this follow-up model was to ensure that cortical atrophy effects were not a major contributing factor to the observed T_2_ relationships with age.

In addition, we conducted a series of paired t-tests (followed by FDR correction of the *p*-values^26^) to examine differences in metabolite T_2_ values between the CSO and PCC voxels. We also computed one linear mixed effects model per metabolite in which we set T_2_ (across both the CSO and PCC voxels) as the outcome variable, age, voxel, and the interaction of age with voxel as the predictors, and a random intercept (u_i_) for each subject: T_2_ = β_0_ + β_1_*(Age-30) + β_2_*Voxel + β_3_*(Age-30)*Voxel + u_i_. The primary aim of this model was to test for any Age*Voxel interaction effects (i.e., whether the age slope differed by brain region in any cases). The linear mixed effects model and random subject intercepts were necessary because this modeling approach structured the data as ‘repeated measures’ in which each participant had two measurements (CSO T_2_ and PCC T_2_).

## 3. Results

### 3.1 Data Quality

Creatine (Cr) linewidths were well within the range of consensus-recommended standards (i.e., < 13 Hz for 3 T^3^) for all spectra except one individual‘s CSO voxel (33-year-old male, Cr linewidth = 14.1 Hz). In addition, for one participant (19-year-old male), the PCC voxel was mistakenly positioned at the wrong location. Thus, these datasets (1 CSO and 1 PCC) were excluded before any statistical analyses. Additional consensus-recommended data quality metrics are presented in Appendix A. Example single-subject spectra at each TE and decay functions for each metabolite are presented in Figure 2. The mean R^2^ value across the whole cohort for the goodness of fit of the T_2_ decay model was ≥0.80 for each of the 6 metabolites of interest and tissue water. Table 2 presents descriptive statistics by age group for the T_2_ values.

**Figure 2.**
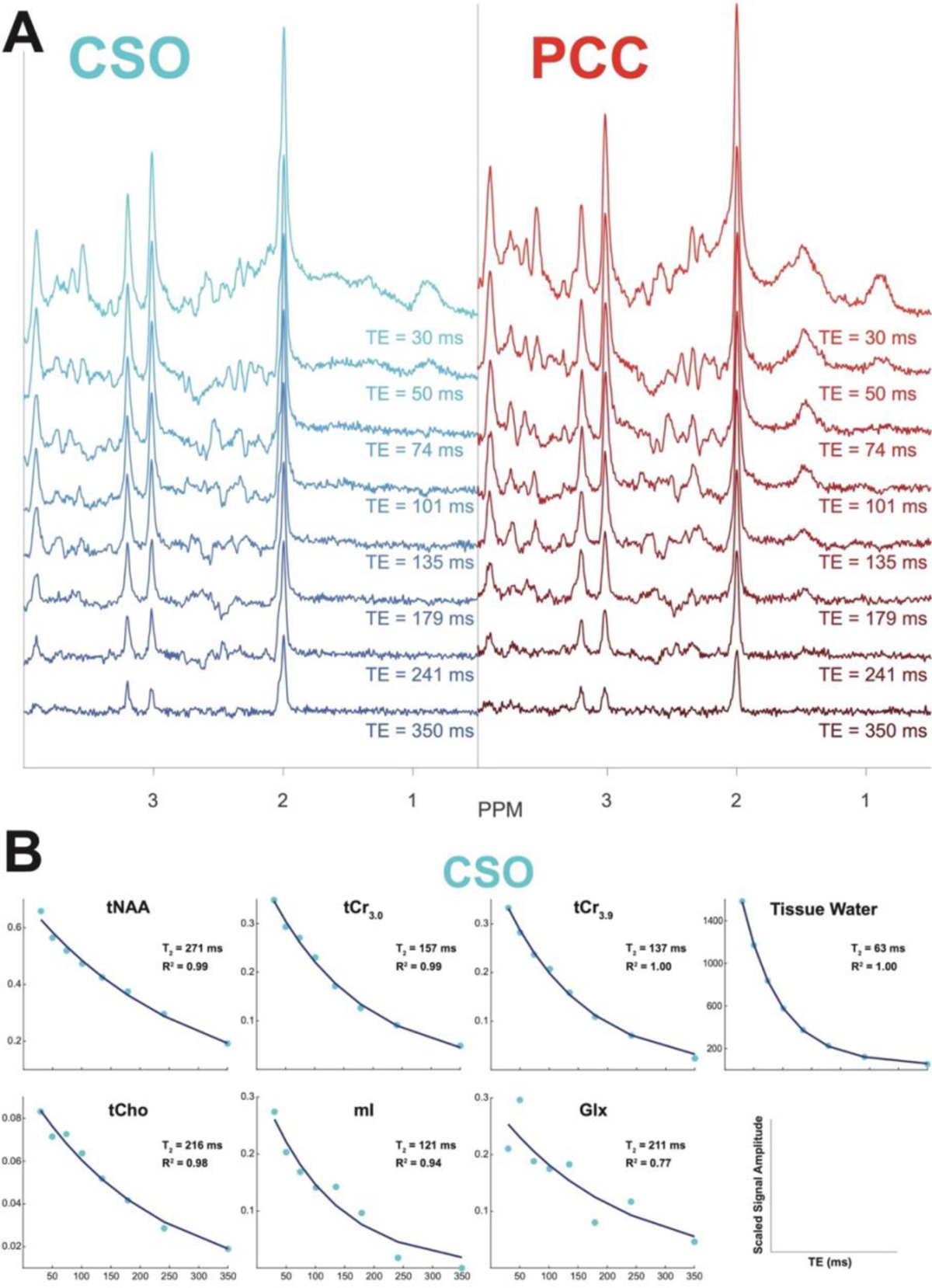
Single-Subject Example Spectra and CSO T2 Decay. A PRESS spectra for each of the 8 TEs for the CSO (left) and PCC (right) voxels for a representative subject (48-year-old female). This representative subject was determined by averaging the R^2^ values of model fit across the 6 metabolites and tissue water for each person in the Johns Hopkins cohort (as water data were unavailable for University of Florida subjects) and then taking the group median of this average R^2^ value. **B** Example T2 decay plots for the 6 metabolites and tissue water from the CSO voxel for the same representative subject. Blue points represent the metabolite amplitude at each TE, and the dark blue line represents the calculated T2 decay curve.

**Table 2.** T_2_ Descriptive Statistics by Age Group.

| **CSO Voxel** |  |  |  |  |  |
| --- | --- | --- | --- | --- | --- |
| **Metabolite** | **18-29 years** | **30-39 years** | **40-49 years** | **50-59 years** | **60+ years** |
|  | Mean (SD) | Mean (SD) | Mean (SD) | Mean (SD) | Mean (SD) |
| tNAA | 298.9<br>(22.5) | 283.8<br>(24.3) | 267.9<br>(24.2) | 267.9<br>(24.6) | 260.6<br>(23.8) |
| tCr<sub>3.0</sub> | 159.5<br>(10.6) | 153.0<br>(8.8) | 147.7<br>(11.5) | 148.3<br>(8.9) | 147.9<br>(8.6) |
| tCr<sub>3.9</sub> | 145.0<br>(10.7) | 133.9<br>(16.2) | 133.0<br>(10.3) | 133.9<br>(8.6) | 132.9<br>(9.3) |
| tCho | 230.1<br>(18.6) | 216.3<br>(24.4) | 203.0<br>(19.7) | 204.5<br>(17.7) | 207.8<br>(19.7) |
| ml | 146.7<br>(14.0) | 141.5<br>(17.2) | 132.7<br>(11.7) | 144.8<br>(16.0) | 140.6<br>(19.0) |
| Glx | 206.1<br>(40.6) | 213.7<br>(36.7) | 186.3<br>(29.6) | 195.6<br>(42.1) | 181.5<br>(30.3) |
| Tissue Water<sup>a</sup> | 64.8<br>(2.1) | 64.1<br>(2.3) | 62.7<br>(2.6) | 62.5<br>(3.1) | 62.4<br>(2.5) |
| **PCC Voxel** |  |  |  |  |  |
| **Metabolite** | **18-29 years** | **30-39 years** | **40-49 years** | **50-59 years** | **60+ years** |
|  | Mean (SD) | Mean (SD) | Mean (SD) | Mean (SD) | Mean (SD) |
| tNAA | 255.2<br>(22.6) | 237.1<br>(12.8) | 224.0<br>(15.3) | 221.1<br>(16.7) | 215.6<br>(14.7) |
| tCr<sub>3.0</sub> | 155.5<br>(8.9) | 149.6<br>(7.8) | 144<br>(8.6) | 145.3<br>(5.8) | 145.8<br>(8.5) |
| tCr<sub>3.9</sub> | 131.1<br>(9.1) | 126.4<br>(5.2) | 125.8<br>(9.6) | 123.6<br>(7.4) | 127.6<br>(10.0) |
| tCho | 258.2<br>(25.2) | 240.3<br>(13.3) | 221<br>(15.9) | 224.2<br>(20.2) | 228.0<br>(20.3) |
| ml | 166.8<br>(15.0) | 157.1<br>(13.7) | 157.5<br>(14.2) | 158.0<br>(13.6) | 159.5<br>(16.3) |
| Glx | 166.8<br>(19.2) | 156.7<br>(21.6) | 164.8<br>(29.3) | 162.9<br>(23.8) | 147.6<br>(31.2) |
| Tissue Water<sup>a</sup> | 54.8<br>(4.0) | 55.2<br>(2.5) | 52.1<br>(2.5) | 49.7<br>(5.9) | 50.7<br>(2.7) |

Note. This table lists T<sub>2</sub> values in ms for the 6 metabolites and tissue water.
<sup>a</sup> Tissue water includes only the $n = 51$ Johns Hopkins subjects.

### 3.2 T_2_ Relationships with Age

Older age was significantly correlated with shorter T_2_ values for tNAA, tCr_3.0_, tCr_3.9_, tCho, Glx, and tissue water in both the CSO and PCC; Spearman r = −0.21 to −0.65, *p* < 0.05, FDR-corrected for multiple comparisons (Figure 3; Table 3). Age was most strongly correlated with tNAA T_2_. Age did not correlate with mI T_2_ for either voxel.

**Figure 3.**
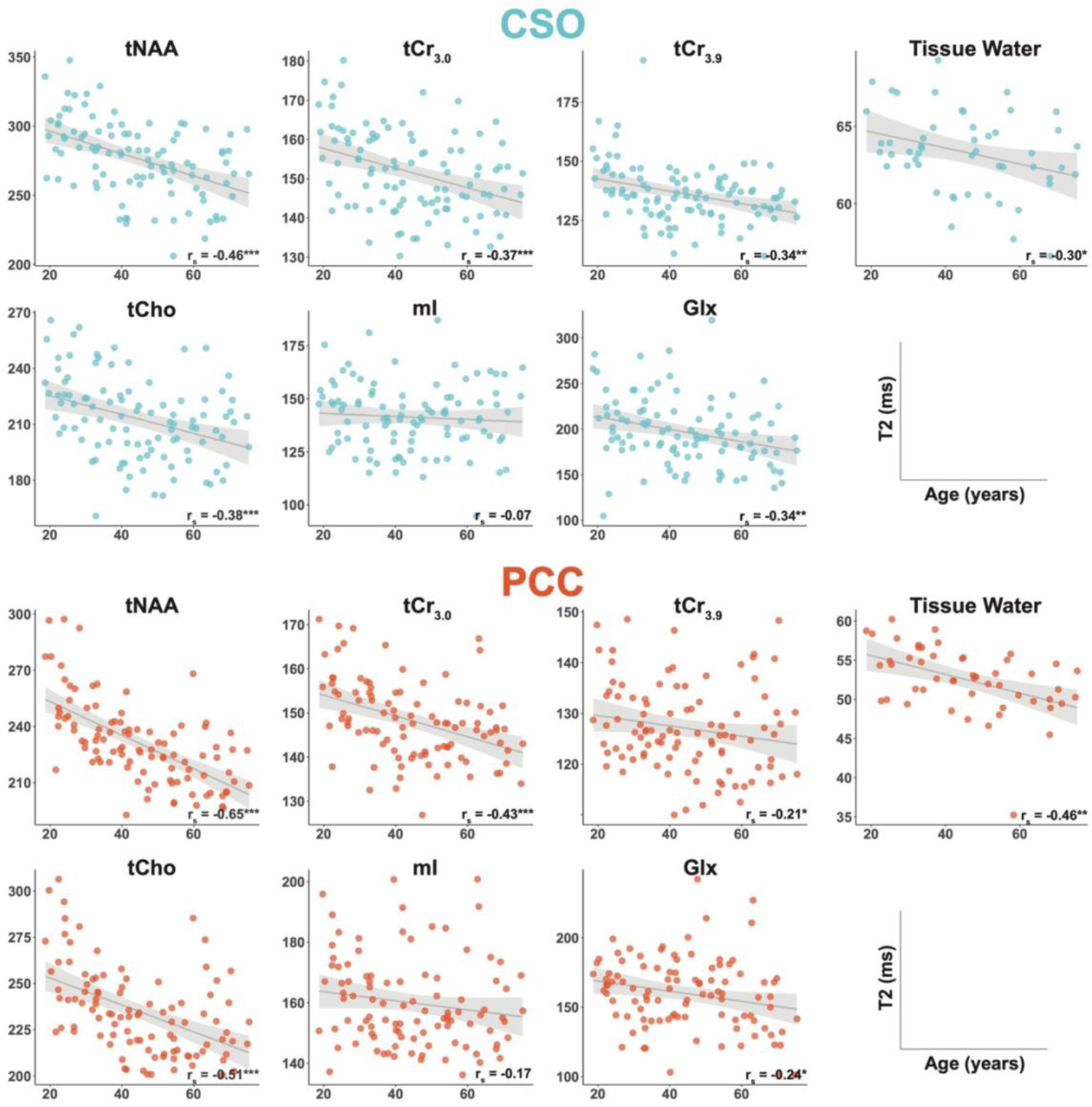
Metabolite T2 Correlations with Age. T2 correlations with age are shown for the CSO (above, blue) and PCC (below, orange). Each point represents one participant. The bottom right corner of each plot indicates the Spearman correlation coefficient (rs) and statistical significance of the FDR-corrected *p*-value for the correlation, \**p*<0.05, \*\**p*<0.01, \*\*\**p*<0.001. The gray line and shading represent the linear model and 95% confidence interval for the model using age to predict T2 (produced using the *geom_smooth* function in R).

**Table 3.**
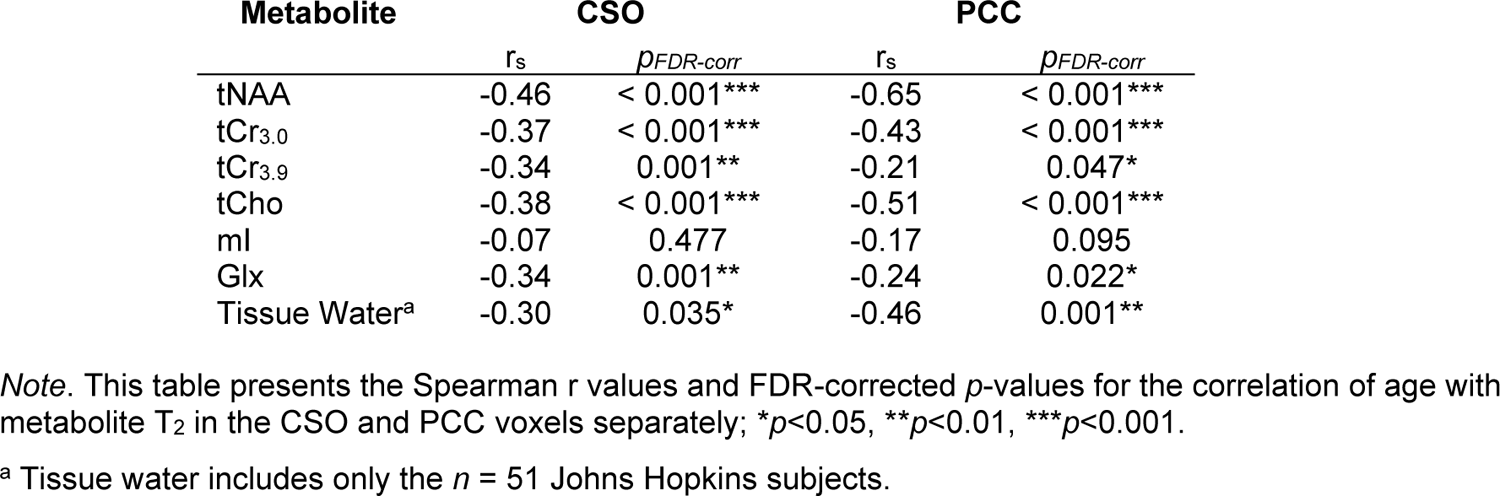
T_2_ Correlations with Age.

Next, we fit linear models to predict T_2_ using age for each metabolite and voxel combination: T_2_ = β_0_ + β_1_*(Age-30); Table 4. As we centered age around 30 years old, the intercept (β_0_) represents the predicted T_2_ value for each metabolite at 30 years of age (as opposed to age 0 which would represent an unhelpful extrapolation). The slope (β_1_) represents the change in the predicted value of T_2_ for each year of life. For example, for tNAA in the CSO, the predicted T_2_ value for an individual age 30 years would be: T_2_ = 288.27 + (30-30)*-0.81 = 288.27 ms, while the predicted T_2_ value for an individual age 50 years would be T_2_ = 288.27 + (50-30)*-0.81 = 272.07 ms. (Note that these predicted T_2_ values also correspond to the gray linear model lines plotted in Figure 3). The slope and intercept values listed in Table 4 can thus be utilized to calculate a predicted T_2_ for any age in a WM- or GM-rich voxel. Table 4 only includes slopes for the metabolites which were significantly correlated with age.

**Table 4.**
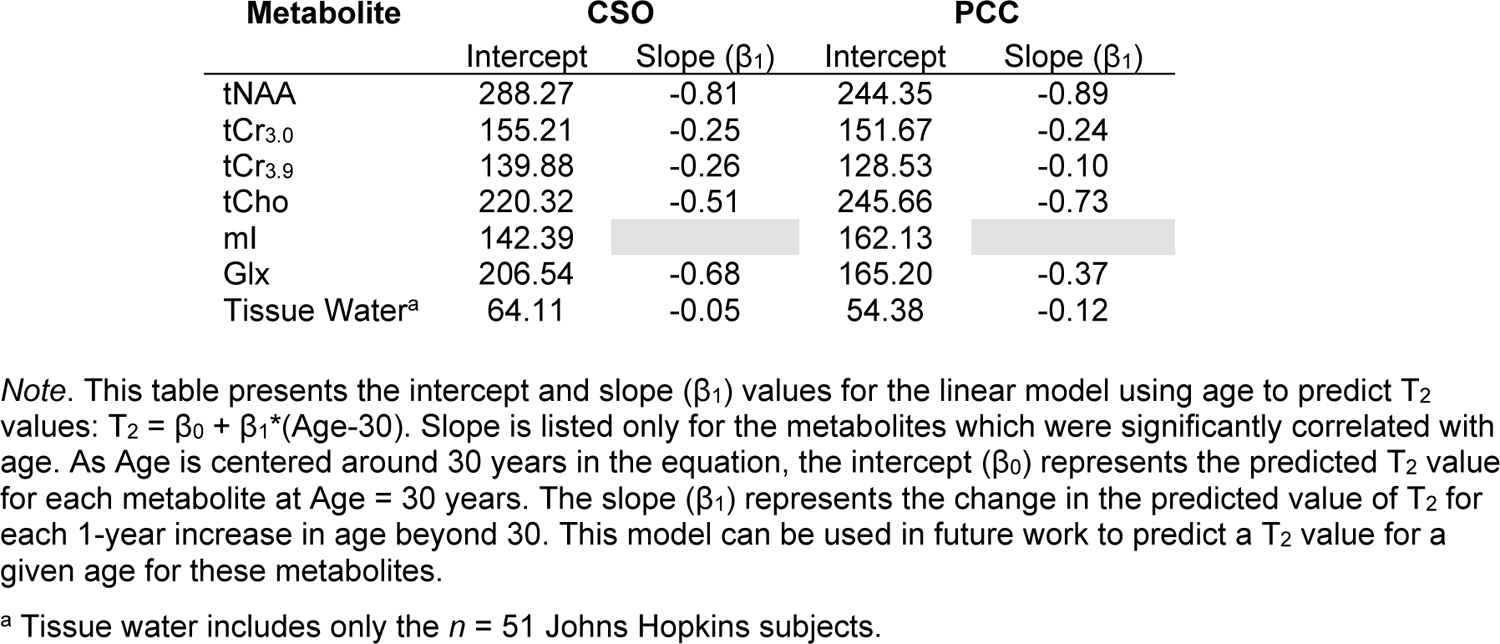
Model Coefficients for Estimating T_2_.

| Metabolite | CSO |  | PCC |  |
| --- | --- | --- | --- | --- |
| | Intercept | Slope ($\beta_1$) | Intercept | Slope ($\beta_1$) |
| tNAA | 288.27 | -0.81 | 244.35 | -0.89 |
| tCr<sub>3.0</sub> | 155.21 | -0.25 | 151.67 | -0.24 |
| tCr<sub>3.9</sub> | 139.88 | -0.26 | 128.53 | -0.10 |
| tCho | 220.32 | -0.51 | 245.66 | -0.73 |
| ml | 142.39 |  | 162.13 |  |
| Glx | 206.54 | -0.68 | 165.20 | -0.37 |
| Tissue Water<sup>a</sup> | 64.11 | -0.05 | 54.38 | -0.12 |

*Note.* This table presents the intercept and slope ( $\beta_1$ ) values for the linear model using age to predict $T_2$ values: $T_2 = \beta_0 + \beta_1(\text{Age}-30)$ . Slope is listed only for the metabolites which were significantly correlated with age. As Age is centered around 30 years in the equation, the intercept ( $\beta_0$ ) represents the predicted $T_2$ value for each metabolite at Age = 30 years. The slope ( $\beta_1$ ) represents the change in the predicted value of $T_2$ for each 1-year increase in age beyond 30. This model can be used in future work to predict a $T_2$ value for a given age for these metabolites.
<sup>a</sup> Tissue water includes only the $n = 51$ Johns Hopkins subjects.

Older age was significantly correlated with greater cortical atrophy (calculated as fGM / (fWM + fGM)) in the PCC (r_s_ = −0.25; *p* = 0.010) but not the CSO (r_s_ = −0.04; *p* = 0.680). As a follow-up to the linear models presented in Table 4, we reran each model controlling for cortical atrophy with aging: T_2_ = β_0_ + β_1_*(Age-30) + β_2_*Tissue (see Supplementary Table B1). Including this metric of cortical atrophy in the model did not change the statistical significance of any T_2_ relationships with age, with the exception of tCr_3.9_ in the PCC (for which the age-T2 relationship became non-significant, *p* = 0.094). Independent of the associations between age and T_2_, greater cortical atrophy was significantly associated with longer metabolite T_2_ values for tNAA, tCr_3.0_, and tCr_3.9_ (CSO only), as well as mI and tissue water (CSO and PCC).

### 3.2 T_2_ Differences by Voxel

Paired t-tests revealed differences in T_2_ values by voxel for all metabolites (as shown in Figure 4). T_2_ values were higher in the CSO than in the PCC for tNAA, tCr_3.0_, tCr_3.9_, Glx, and tissue water, and lower in the CSO for tCho and mI.

**Figure 4.**
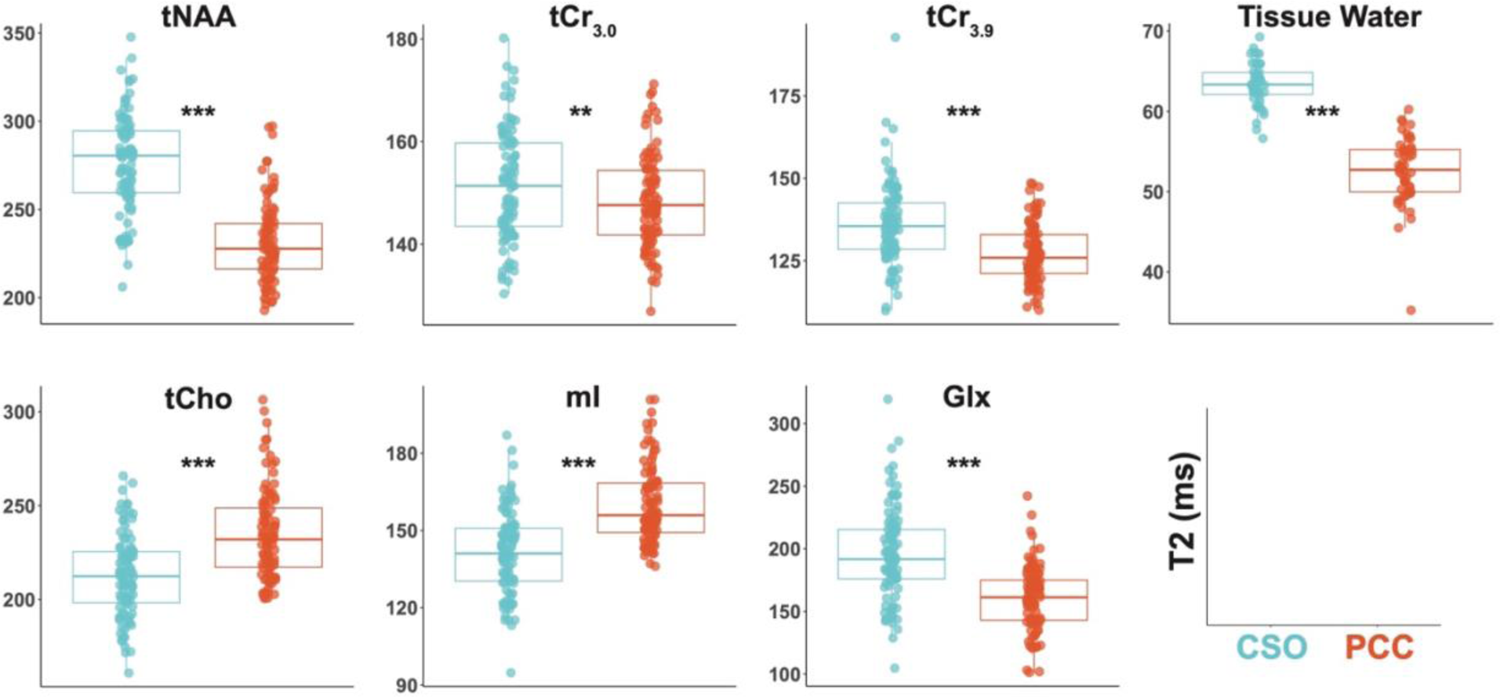
Metabolite T2 Differences by Voxel. T2 differences by voxel are shown for the CSO (blue) and PCC (orange). Each point represents one participant. The asterisks indicate the statistical significance of the FDR-corrected *p*-value for the paired t-test, \**p*<0.05, \*\**p*<0.01, \*\*\**p*<0.001.

As a follow-up analysis, we computed one linear mixed effects model per metabolite (across both the CSO and PCC) to test whether the age slope differed by brain region for any metabolites or tissue water. The Age*Voxel interaction was significant only for tissue water (*p*=0.007), indicating that, in all but one case, the relationship of age with T_2_ did not differ based on brain region (Supplementary Table B2). However, for tissue water, the relationship of age with T_2_ was stronger for the PCC than the CSO.

## 4. Discussion

Here we present the largest analysis to date examining metabolite and tissue water T_2_ changes across the healthy adult lifespan. Among 101 adults ages 18-75 and across two sites, metabolite and tissue water T_2_ values in both the CSO and PCC were generally significantly shortened with age, even when controlling for age-related cortical atrophy. Moreover, T_2_ values were longer in the CSO, with the exception of tCho and mI which exhibited longer T_2_ in the PCC. Taken together, these results align with the majority of prior work which also reported T_2_ declines with normal aging (but in much smaller cohorts). Moreover, the finding of T_2_ differences based on age and brain region highlights the importance of measuring subject-level T_2_ during data acquisition or employing estimation methods (such as the statistical models provided here) for calculating age- and region-appropriate T_2_ values.

Older age was correlated with shorter metabolite T_2_ values for tNAA, tCr_3.0_, tCr_3.9_, tCho, Glx, and tissue water in both the CSO and PCC. This aligns with most prior research in smaller samples which similarly found shorter tNAA, tCr, tCho, and tissue water T_2_s with older age^4–7^. As also seen in these prior studies, we identified the strongest age association for tNAA. Most prior reports did not examine Glx. Deelchand and colleagues (2020) reported reduced mI T_2_ in CSO and PCC in older age, whereas we did not find an association between mI T_2_ and age. However, it should be noted that Deelchand and colleagues^4^ compared two age groups (rather than treating age as a continuous variable), and their older cohort (ages 70-83 years) extended beyond our upper age range.

The specific mechanisms underlying these observed T_2_ changes with aging remain unclear. With the exception of tCr_3.9_ in the PCC, each of the identified T_2_ associations with age remained statistically significant when controlling for cortical atrophy, suggesting that age-related atrophy is not a major factor in these findings. Instead, as metabolites are largely intracellular (glial or neuronal)^28^, their T_2_ relaxation times are likely influenced by changes in the cellular microenvironment^4^, i.e. cellular morphology, metabolism, or myelination^4,7^. It is well established that neurons undergo morphological changes during aging—such as reduction in soma size and loss or regression of dendrites and dendritic spines^29^— alongside a parallel metabolic shift in astrocytes associated with increased neuroinflammatory response and changes in oxidative metabolism^30,31^. Furthermore, degeneration of myelin sheath^32^ and loss of axonal fiber^32^ with advancing age, accompanied by debris (e.g., protein aggregates) and degraded myelin accumulation^33,34^ reported in white matter and further supported by *in vivo* diffusion tensor imaging^35,36^.

The observed T_2_ changes could also be influenced by the gradual deposition of iron, particularly Fe^3+^, in the brain with aging. Although iron is present in the brain in multiple forms, the intracellular non-heme iron (i.e., ferritin) in tissue is thought to cause dephasing of the proton spins and thus a faster T_2_ decay^37^. Several studies observed a strong linear correlation between iron concentrations and transverse relaxation (R_2_)^38^ values both *in vivo* and in post-mortem healthy and Alzheimer’s disease brain tissue^37^, suggesting that faster T_2_ relaxation is related to age-related iron deposition^39^. Whilst the precise contribution of each of these mechanisms is unclear, the observed age relationships suggest that T_2_ measurements are sensitive to various parallel changes in the cellular environment^7^.

In this dataset, T_2_ relaxation times were predominantly longer for tNAA, tCr_3.0_, tCr_3.9_, Glx, and tissue water in the WM-rich CSO, whereas tCho and mI exhibited longer T_2_ in the GM-rich PCC. There is relatively limited literature considering GM/WM differences in metabolite T_2_, and this prior work differs in voxel location, cohort, acquisition and quantification methodology, and statistical approach. Of the seven references we identified^6,40–45^, five reported longer T_2_ for NAA in WM as we did^6,40–42,45^, while one revealed no significant tissue effect^43^ and one showed the reverse effect^44^. For tCr_3.0_, three references found our result of longer T_2_ in WM^6,41,42^, three showed no difference^40,43,45^, and the same study showed the reverse effect^44^. Only a few studies have measured T_2_s of tCr_3.9_ or Glx. For tCr_3.9_, one paper showed longer T_2_s in WM^45^ (as we found) and one no difference^42^; for Glx, one study that separated Glx as Glu and Gln with J-PRESS found no differences by tissue type in the major component, Glu^42^. For tCho, four studies found no difference^40,41,43,45^, two found longer T_2_ in GM^42,44^ (as we did), and one longer T_2_ in WM^6^. T_2_ of mI remains less investigated, but the two studies that measured mI T_2_ also found longer T_2_ in GM^42,44^. For tissue water, GM is generally found to have longer T_2_ values than WM in multi-echo MRI experiments^46–48^, although the extent to which CSF confounds this result depends on resolution. Regional T_2_ differences may relate to greater micro- and macro-structural organization in myelinated WM compared to GM^49^; however, further work is needed to fully understand the mechanisms that govern metabolite T_2_ relaxation.

We recently performed a meta-regression analysis of 75 manuscripts^50^ containing 629 unique values to derive a general predictive T_2_ model, with linear factors for: metabolite, field strength, species, tissue, pulse sequence, and Carr-Purcell Meiboom-Gill filter. The average bias between the (30-year-old intercept) values reported in the present study and the model predicted values was +9 ms (i.e., on average the model predicts shorter T_2_s than measured here). The average absolute difference was 23 ms which is smaller than the average absolute difference between the predicted model and the T_2_ training dataset (42 ms). On this basis, we assert that our results are consistent with the diverse T_2_ literature.

2D modeling of interrelated MRS data has recently gained interest in the MRS community^51–53^. Most notably, it was found that 2D modeling of synthetic multi-TE MRS data with overlapping peaks led to improved precision due to improved model parsimony achieved through reparametrization^51^. Applying 2D modeling to our *in vivo* datasets may improve the T_2_ estimation of the metabolites reported here and could potentially allow for the T_2_ estimation of additional low-SNR metabolites. However, it will also require careful reparameterization of the T_2_ relaxation constants, lineshape estimates, and baseline terms, which will be part of future studies.

There are several limitations to this work. First, we acknowledge that our T_2_ measurement here is a complex mix of pure T_2_ and some inhomogeneous broadening factors that are not fully refocused by the two PRESS spin echoes. The goal of the present work was to improve the accuracy of T_2_ relaxation correction in quantification procedures by understanding age effects on our measure of T_2_ (rather than to accurately measure pure T_2_). Second, future work could expand upon the age range to include those younger than 18 and older than 70 years, as well as targeting both normal and pathological aging (e.g., Alzheimer’s and other neurodegenerative diseases). Given the potential of T_2_ to reflect both micro- and macrostructural organization, the measure may show utility as an early indicator of these changes, as suggested by Kirov and Tal^54^. Though this was a large cohort with systematic recruitment across the adult lifespan, we only enrolled a few individuals older than age 70 years (the timeframe at which aging effects drastically accelerate). Lastly, we were limited to collecting only two voxels (WM-rich CSO and GM-rich PCC); however, prior evidence suggests that neurochemical changes with aging are highly region-dependent^55^, and therefore future work might consider probing T_2_ changes in other brain regions, or across the entire brain.

## 5. Conclusions

Consistent with prior literature, in a large multi-site cohort sampled systematically across the adult lifespan, we identified a clear age-related decrease in T_2_ for multiple metabolites and tissue water, as well as differences in T_2_ between the WM-rich CSO and GM-rich PCC. Together, these findings highlight potential changes in the brain’s cellular microenvironment with normal aging and underscore the critical importance of considering metabolite T_2_ differences across the adult lifespan in MRS quantification procedures. We suggest that future MRS work leverage the models presented here to estimate age- and region-specific T_2_ values instead of relying on uniform default values.

## Author Contributions

KH processed all data, conducted all statistical analyses, prepared all figures and supplemental material, and prepared the manuscript. SM contributed to protocol development, manuscript writing and led all revisions of the manuscript. HZ contributed to MRS data processing, and developed Osprey code for the analysis. YS and EC made significant contributions to data collection. CDJ generated the spectra figure and contributed to interpretation of results. AG, DS, and GS contributed to interpretation of results and drafted parts of the Discussion. VY reviewed all structural scans to assess data quality and check for incidental findings. SH set up the scan protocol and oversaw data quality control. GO, EP, and RAE designed the project and led interpretation of the results. All authors participated in revision of the manuscript.

## Acknowledgements

The authors also wish to thank all of the participants who volunteered their time, as well as support staff at both MRI centers, without whom this project would not have been possible.

