## Appendix A for "Metabolite T_2_ relaxation times decrease across the adult lifespan in a large multi-site cohort"

### Appendix A: Summary MRSinMRS Report

Here we provide a summary following the minimum reporting standards in MRS generated in Osprey. For further details, see: Lin et al. 'Minimum Reporting Standards for *in vivo* Magnetic Resonance Spectroscopy (MRSinMRS): Experts' consensus recommendations. NMR in Biomedicine. 2021;e4484. doi.org/10.1002/nbm.4448

**Sites:** The Johns Hopkins University School of Medicine / F. M. Kirby Research Center for Functional Brain Imaging, Kennedy Krieger Institute and the University of Florida McKnight Brain Institute.

#### 1. Hardware

**Johns Hopkins University**

| | |
|---|---|
| a. Field strength [T] | 3 T |
| b. Manufacturer | Philips |
| c. Model (software version if available) | R 5.7.1 |
| d. RF coils: nuclei (transmit/receive), number of channels, type, body part | 1H, 32 channel, head |

**University of Florida**

| | |
|---|---|
| a. Field strength [T] | 3 T |
| b. Manufacturer | Philips |
| c. Model (software version if available) | R 5.9.0 |
| d. RF coils: nuclei (transmit/receive), number of channels, type, body part | 1H, 32 channel, head |

#### 2. Acquisition

**Metabolite TE Series**

| | |
|---|---|
| a. Pulse sequence | Philips PRESS |
| b. Volume of interest (VOI) locations | Right CSO, Midline PCC |
| c. Nominal VOI size [$mm^3$] | 30 x 26 x 26 $mm^3$ |
| d. Repetition time (TR), echo time (TE) [ms] | TR 2000 ms,<br>TE 30, 50, 74, 101, 135, 179, 241, 350 ms |
| e. Total number of averages per spectrum | 24 averages per TE = 192 averages in total |
| f. Additional sequence parameters | F1: 2000 Hz, 1024 points |
| g. Water suppression method | CHESS (115 Hz bandwidth) |
| h. Shimming method, reference peak | 2nd order pencil beam, water |
| i. Trigger or motion correction | No trigger or active motion correction |

**Water TE Series**

| | |
|---|---|
| a. Pulse sequence | Philips PRESS |
| b. Volume of interest (VOI) locations | Right CSO, Midline PCC |
| c. Nominal VOI size [$mm^3$] | 30 x 26 x 26 $mm^3$ |
| d. Repetition time (TR), echo time (TE) [ms] | TR 2000 ms,<br>TE 30, 50, 74, 101, 135, 179, 241, 350 ms |
| e. Total number of averages per spectrum | 2 averages per TE = 16 averages in total |
| f. Additional sequence parameters | F1: 2000 Hz, 1024 points |
| g. Water suppression method | None |
| h. Shimming method, reference peak | 2nd order pencil beam, water |
| i. Trigger or motion correction | No trigger or active motion correction |

| 3. Data analysis methods and outputs | |
|---|---|
| a. Analysis software | Osprey 2.5.0 |
| b. Processing steps deviating from Osprey | None |
| c. Output measure | Metabolite amplitude (from A_amplMets_Voxel_1_Basis_1.tsv) |
| d. Quantification references and assumptions, fitting model assumptions | Basis set list: Asc,Asp,Cr,-CrCH2,GABA,GPC,GSH, Gln,Glu,Lac,mI,NAA,NAAG,PCh,PCr,PE,sI,Tau, MM09,MM12,MM14,MM17,MM20,Lip09,Lip13,Lip20<br>Fitting method: Osprey baseline knot spacing 0.40 ppm |

| 4. Data quality | |
|---|---|
| **Metabolite TE Series, CSO** | |
| a. SNR (Cr), linewidth (Cr) [Hz] | SNR: 86 ± 15, linewidth: 7.01 ± 1.12 Hz |
| b. Data exclusion criteria | Cr linewidth > 13 Hz |
| c. Quality measures of postprocessing model fitting (Mean Relative Amplitude Residual) | 5.40 % |
| **Water TE Series, CSO** | |
| a. SNR (water), linewidth (water) [Hz] | SNR: 565 ± 106, linewidth: 7.13 ± 0.86 Hz |
| b. Data exclusion criteria | None |
| **Metabolite TE Series, PCC** | |
| a. SNR (Cr), linewidth (Cr) [Hz] | SNR: 89 ± 19, linewidth: 6.23 ± 0.88 Hz |
| b. Data exclusion criteria | Cr linewidth > 13 Hz |
| c. Quality measures of postprocessing model fitting (Mean Relative Amplitude Residual) | 6.79 % |
| **Water TE Series, PCC** | |
| a. SNR (water), linewidth (water) [Hz] | SNR: 529 ± 118, linewidth: 6.59 ± 0.77 Hz |
| b. Data exclusion criteria | None |

*Note.* SNR, linewidth, and relative amplitude residual metrics are reported for TE: 30 ms only. Cr SNR is the maximum amplitude of the tCr peak divided by the detrended standard deviation of the noise. Mean relative amplitude residual is indicative of postprocessing model fit (in which higher values indicate poorer model fit).
