## Supplementary Table B1 for "Metabolite T_2_ relaxation times decrease across the adult lifespan in a large multi-site cohort"

### **Appendix B.** Supplemental Tables

#### **Table B1.** Model Results Including Cortical Atrophy

| **Metabolite** | **CSO** | | **PCC** | |
|---|---|---|---|---|
| | Age *p*-value | Tissue *p*-value | Age *p*-value | Tissue *p*-value |
| tNAA | < 0.001*** | < 0.001*** | < 0.001*** | 0.350 |
| $tCr_{3.0}$ | < 0.001*** | < 0.001*** | < 0.001*** | 0.102 |
| $tCr_{3.9}$ | < 0.001*** | 0.015* | 0.094 | 0.781 |
| tCho | < 0.001*** | 0.085 | < 0.001*** | 0.359 |
| mI | 0.403 | 0.013* | 0.436 | 0.003** |
| Glx | 0.003** | 0.188 | 0.009** | 0.078 |
| Tissue Water[a] | 0.045* | 0.028* | < 0.001*** | < 0.001*** |

*Note*. This table presents the *p*-values for the linear models using age and cortical atrophy (i.e., the relative tissue fraction within the voxel, fGM / (fWM + fGM)) to predict metabolite $T_2$ value: $T_2 = \beta_0 + \beta_1$*(Age-30) + $\beta_2$*Tissue. *$p<0.05$, **$p<0.01$, ***$p<0.001$

[a] Tissue water includes only the $n$ = 51 Johns Hopkins subjects.

#### **Table B2.** Linear Mixed Effects Model Results Including Age*Voxel Interaction

| **Metabolite** | **Age *p*-value** | **Voxel *p*-value** | **Age*Voxel *p*-value** |
|---|---|---|---|
| tNAA | <0.001*** | <0.001*** | 0.769 |
| $tCr_{3.0}$ | <0.001*** | 0.222 | 0.854 |
| $tCr_{3.9}$ | <0.001*** | <0.001*** | 0.066 |
| tCho | 0.001** | <0.001*** | 0.104 |
| mI | 0.549 | <0.001*** | 0.473 |
| Glx | 0.003** | 0.001** | 0.443 |
| Tissue Water[a] | 0.073 | <0.001*** | 0.007** |

*Note*. This table presents the *p*-values for the linear mixed effects models using Age, Voxel, and the interaction of Age*Voxel, as well as a random intercept for each subject ($u_i$), to predict metabolite $T_2$ value: $T_2 = \beta_0 + \beta_1$*(Age-30) + $\beta_2$*Voxel + $\beta_3$*(Age-30)*Voxel + $u_i$. *$p<0.05$, **$p<0.01$, ***$p<0.001$.

[a] Tissue water includes only the $n$ = 51 Johns Hopkins subjects.
